# Microbiota-derived metabolites inhibit *Salmonella* virulent subpopulation development by acting on single-cell behaviors

**DOI:** 10.1101/2021.02.14.430798

**Authors:** Alyson M Hockenberry, Gabriele Micali, Gabriella Takács, Jessica Weng, Wolf-Dietrich Hardt, Martin Ackermann

**Affiliations:** Department of Environmental Systems Science, ETH Zürich, Zürich, 8092, Switerland; Department of Environmental Microbiology, Eawag, Dübendorf, 8600, Switzerland; Mayo Clinic Alix School of Medicine, Mayo Clinic College of Medicine and Science, Mayo Clinic, Rochester, Minnesota, 55905, U.S.A; Institute of Microbiology, D-BIOL, ETH Zürich, Zürich, 8092, Switzerland

## Abstract

*Salmonella spp.* express *Salmonella* pathogenicity island 1 (SPI-1) genes to mediate the initial phase of interaction with host cells. Prior studies indicate short-chain fatty acids, microbial metabolites at high concentrations in the gastrointestinal tract, limit SPI-1 gene expression. A number of reports show only a subset of *Salmonella* cells in a population express these genes, suggesting short-chain fatty acids could decrease SPI-1 population-level expression by acting on per-cell expression and/or the proportion of expressing cells. Here, we combine single-cell, theoretical, and molecular approaches to address the effect of short-chain fatty acids on SPI-1 expression. Our results show short-chain fatty acids do not repress SPI-1 expression by individual cells. Rather, these compounds act to selectively slow the growth of SPI-1 expressing cells, ultimately decreasing their frequency in the population. Further experiments indicate slowed growth arises from short-chain fatty acid-mediated depletion of the proton motive force. By influencing the SPI-1 cell-type proportions, our findings imply gut microbial metabolites act on cooperation between the two cell-types and ultimately influence *Salmonella*’s capacity to establish within a host.

**Significance Statement:** Emergence of distinct cell-types in populations of genetically identical bacteria is common. Furthermore, it is becoming increasingly clear that cooperation between cell-types can be beneficial. This is the case during *Salmonella* infection, in which cooperation between inflammation-inducing virulent and fast-growing avirulent cell-types occurs during infection to aid in colonization of the host gut. Here, we show gut microbiota-derived metabolites slow growth by the virulent cell-type. Our study implies microbial metabolites shape cooperative interactions between the virulent and avirulent cell types, a finding that can help explain the wide array of clinical manifestations of *Salmonella* infection.

## Introduction

The mammalian gastrointestinal tract (GI) is chemically defined by resident bacteria metabolizing the host’s diet. Among the most abundant microbial metabolites are the short chain fatty acids (SCFAs) acetate, butyrate, and propionate. Each are found at up to 100 millimolar concentrations in the human GI, with levels lowest in the ileum, increasing to high levels in the proximal colon, and tapering off in the distal colon (1). They also vary as a function of microbiota members and fluctuate over time, correlating with the timing and composition of meals (2–4). The chemical environment an enteric microbe finds itself in, therefore, varies by location in the gut, from person-to-person, and across time.

This holds true as well during the initial phases of infection by the enteric pathogen *Salmonella enterica. Salmonella* expresses virulence genes to inflame and subsequently colonize the host GI (reviewed in (5)). Inflammation is initially caused by invading into gut epithelial tissues using a Type III Secretion System encoded on *Salmonella* pathogenicity island 1 (SPI-1, (6, reviewed in 7)). This secretion system pumps effector proteins directly into host cells, leading to bacterial uptake and subsequent inflammation. Inflammation increases nutrient availability and killing of competitor resident microbiota, opening a niche for *Salmonella* establishment in the gut (8–12).

Recent evidence shows there is cell-to-cell variation in SPI-1 expression by *Salmonella* cells. Each individual in the population is found in a discrete expressing (SPI-1+) or non-expressing (SPI-1−) state (13, 14). The emergence of these two cell-types is beneficial as they cooperate with each other to facilitate colonization of the host. SPI-1+ cells invade host tissues and induce inflammation while SPI-1− cells replicate quickly in the intestinal lumen to exploit the niche cleared by the SPI-1+ cells (15–17). It is currently unclear whether these cooperative interactions are influenced by environmental signals; of particular interest are the dynamic conditions of the GI, where there is high variation in the frequency of SPI-1+ cells (18).

Several reports have shown microbiota-derived metabolites impact *Salmonella* infectivity. Importantly, SCFAs at physiological levels decrease SPI-1 expression by populations of *Salmonella* cells and limit their invasion into host cells (19, 20). The authors concluded from these studies that SCFAs repress SPI-1 expression and result in decreased pathogenicity. Consistent with this idea, the SCFAs butyrate and propionate impact *Salmonella* pathogenicity *in vivo* (21, 22). However, because SPI-1 expression varies from cell-to-cell within a population prompts revisiting the effect of SCFAs on SPI-1 expression from a single-cell perspective.

The previously observed SCFA-mediated reduction in population-level SPI-1 expression could arise from a number of mechanisms scaling from behaviors by individual cells. Determining how SCFAs impact single-cell behaviors will inform on their mechanism of action and ultimately how the gut environment influences *Salmonella* pathogenicity.

## Results + Discussion

### SCFAs repress population-level SPI-1 expression through decreasing the proportion of SPI-1+ cells

Reduced population-level SPI-1 expression could result from a combination of mechanisms: either by fewer SPI-1+ cells in the population or by decreased SPI-1 expression per-cell. To address how SCFAs impact SPI-1 expression by individual cells during growth, we first quantified population-level growth and SPI-1 expression using SPI-1 transcriptional reporter cells (*PprgH-gfp*) grown in a range of SCFA concentrations (Figs 1A,B, Table S1). Consistent with previous reports (19, 20), increasing SCFA concentration correlated with decreased population-level SPI-1 expression (Fig 1C, *one-way anova*, *p < 0.0001*). Increased SCFA concentrations also increased population-level lag time and maximal growth rate (Fig S1, *one-way anovas*, *p < 0.0001, p < 0.005, respectively*).

**Fig 1:**
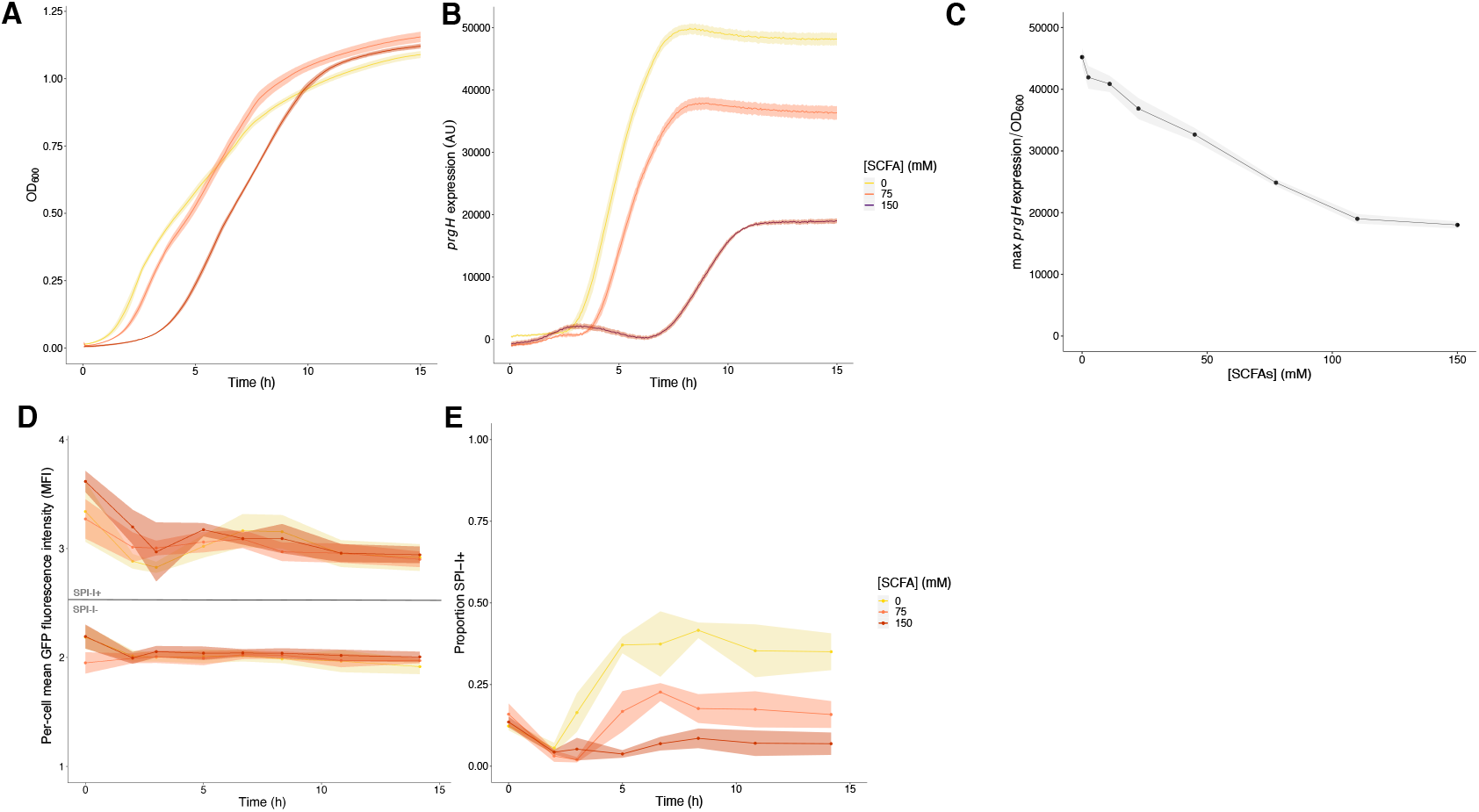
SCFAs decrease population-level SPI-1 expression by decreasing the proportion of SPI-1+ cells. **A** Mean ± standard error of the mean (SEM) OD600 and **B** mean ± SEM GFP fluorescence intensity (*i.e.* SPI-1 expression) by SPI-1 reporter cells in no (0 mM, yellow), intermediate (75 mM, salmon), and high (150 mM, dark orange) SCFA concentrations measured every 3 min over 15 h in a plate reader assay **C** SPI-1 expression normalized to OD600 as a function of SCFA concentration. Average of triplicates from 3 independent experiments, one-way ANOVA, expression vs SCFA concentration, p < 0.0001; Tukey’s Honest Significance Test, 0 mM vs 75 mM, p < 0.0001, 75 mM vs 150 mM, p < 0.001. The impact of SCFAs on single-cell **D** mean GFP fluorescence intensity (MFI) ± SEM by SPI-1 cell-type and **E** mean ± SEM proportion of SPI-1+ cells as measured by flow cytometry. 50,000 cells quantified per timepoint and averaged 3 independent experiments; MFI by treatment over time, two-way ANOVA, MFI by SCFA treatment, p > 0.2; SPI-1 cell type proportions by SCFA treatment over time, two-way ANOVA, proportion by SCFA treatment, p < 0.0001.

Using intermediate (75 mM) or high (150 mM) levels of SCFAs shown to reduce SPI-1 expression, we next quantified SPI-1 expression on a per-cell basis. Reporter cells were cultured with the 0, 75, or 150 mM SCFAs and examined by flow cytometry at a range of timepoints. The mean fluorescence intensity of individual SPI-1+ cells was unaffected by SCFA treatment (Fig 1D, *two-way anova: SPI-1+ vs SPI-1−, p < 0.001; SCFA treatment, p > 0.25*). Rather, SCFA treatment resulted in a dose-dependent decrease in the proportion of SPI-1+ cells (Fig 1E, *two-way anova*, *p < 0.0001*). Untreated populations of cells consisted of roughly 40% SPI-1+ cells by mid-exponential phase through stationary phase. Populations grown with moderate or high SCFA concentrations consisted of 25% and 10% SPI-1+ cells at these timepoints, respectively. These observations demonstrate SCFAs do not decrease SPI-1 expression by individual cells, rather they act on the frequency of SPI-1+ cells in the population.

### The SCFA-mediated decrease in SPI-1+ cells occurs through limitations on SPI-1+ cell growth

How can SCFAs decrease the frequency of SPI-1+ cells in the population? We reasoned SPI-1+ population frequency is set by four parameters: the growth rates of SPI-1- and SPI-1+ cells and the rates of switching between the two cell-types (Fig 2A). A minimal mathematical model (expanded up in *Supporting Materials*) established decreased SPI-1+ frequency can arise through multiple, non-mutually exclusive scenarios: increasing the SPI-1+ to SPI-1− switching rate, decreasing the SPI-1- to SPI-1+ switching rate, increasing the SPI-1− growth rate, or decreasing the SPI-1+ growth rate (Figs 2B,C,D). Parameter space exploration shows the interaction of these four parameters (Figs 2C, D). In general, smaller changes in growth rates led to large shifts in the cell-type proportions, whereas larger changes in switching rates were necessary to shift cell-type proportions.

**Fig 2:**
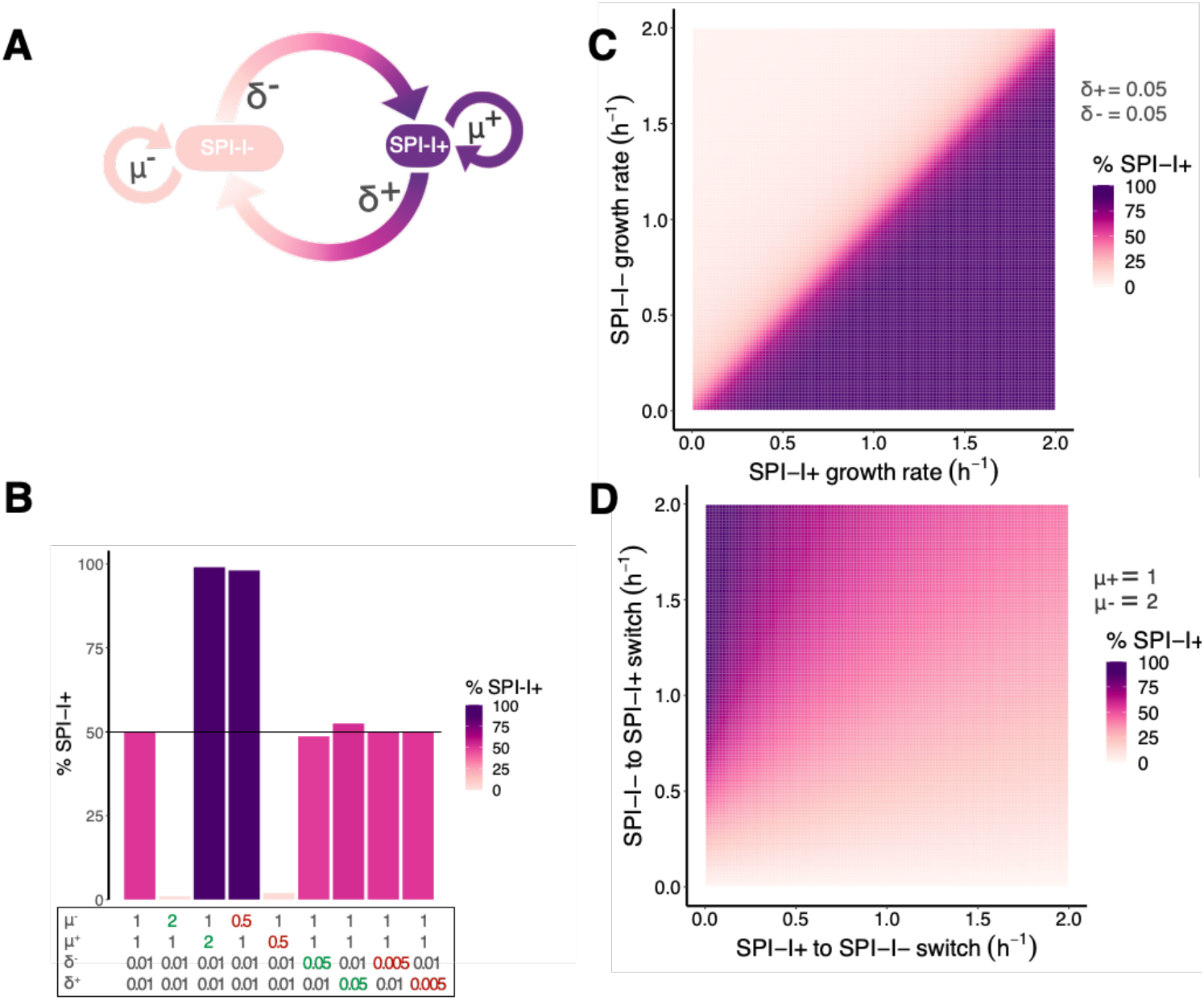
Mathematical modelling shows single-cell growth and cell-type switching rates shape subpopulation frequencies. **A** Conceptual model underlying SPI-1 cell-type frequencies. The cell-type frequency is set by these four parameters: μ−, growth rate of SPI-1− cells; δ−, SPI-1− to SPI-1+ switching rate; μ+, growth rate of SPI-1+ cells; and δ+, SPI-1− to SPI-1+ switching rate. **B** The conceptual model from panel A was transformed into mathematical equations to estimate how these parameters impact cell-type frequency analytically (expanded upon in the *Supplementary Materials*). Calculated SPI-1+ proportions assuming the indicated rates of growth and cell-type switching. **C** How SPI-1 cell-type growth rates impact SPI-1+ proportion when δ+ = 0.05 h^−1^ and δ− = 0.05 h^−1^ **D** How SPI-1 cell-type switching rates impact SPI-1+ proportion when μ− = 2 h^−1^and μ+ = 1 h^−1^

Knowing how SCFAs change these parameters inform on their mechanism of action. Changes in cell-type growth rates imply SCFAs act on cellular metabolism. Meanwhile, SCFA action on cell-type switching rates would suggest they act on SPI-1 regulation by individual cells. We measured impact of SCFAs on these four single-cell parameters directly using quantitative time-lapse microscopy. We employed a “feeding culture” microfluidic approach to recapitulate our population-level experiments (23). SPI-1 reporter cells under observation in a microfluidic chip were fed by an actively growing *Salmonella* culture with or without SCFAs. In this manner, reporter cells under observation are experiencing the dynamic environment of a growing culture, where nutrients are depleted and compounds are excreted. We observed individual reporter cells over 12 h of culture (Fig 3A, Supp video 1). By analyzing the images, we quantified single-cell growth rates and cell-type switches for each cell-type across growth phases.

**Fig 3:**
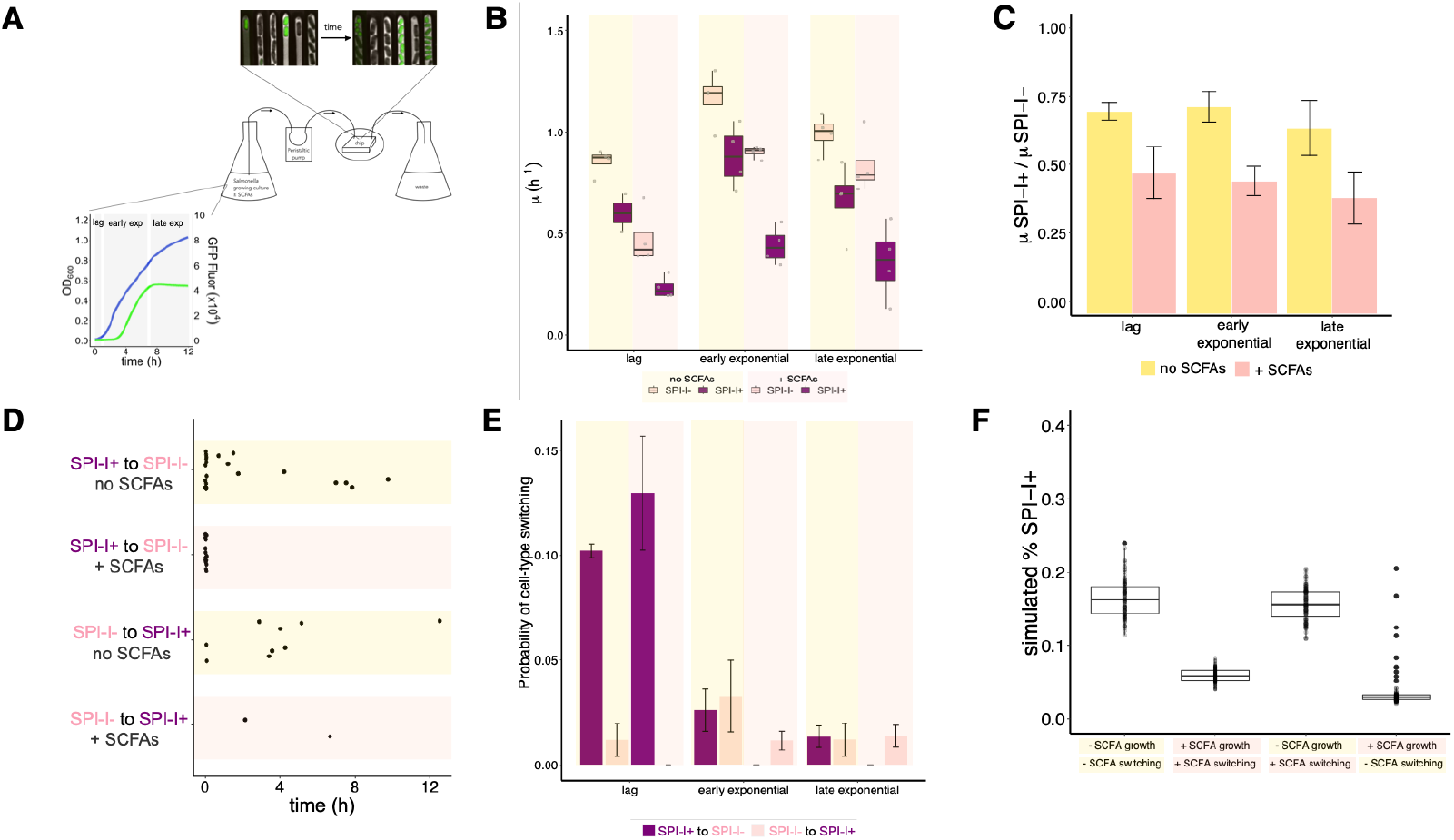
SCFAs slow growth and inhibit phenotypic switching by *Salmonella* cells. **A** Experimental set-up for quantitative time-lapse microscopy using a feeding culture approach. An actively growing culture of *Salmonella* cells is pumped through a microfluidic chip and into a waste container using a peristaltic pump. Simultaneously, SPI-1 reporter *Salmonella* cells loaded into the microfluidic chip are experiencing the same chemical environment as cells in the flask. Cells in the chip are imaged every 3 minutes. Time-lapse images are then analyzed to extract SPI-1− and SPI-1+ growth and switching rates. Data presented is from 4 independent experiments; we designated the first 0.5 h of growth lag phase, the next 6 h early exponential, and the last 5.5 h late exponential. **B** Mean growth rates (n = 2502 cells, box = quartiles, whiskers = all observations) by cell-type, SCFA treatment, and growth phase. Three-way ANOVA followed by Tukey’s Honest Significance Test, ** p < 0.001 **C** Ratio ± SEM of SPI-1+ and SPI-1− cell growth rates in the absence of presence of SCFAs. Two-way ANOVA followed by Tukey’s Honest Significance test, ** p < 0.001. **D** Observed cell-type switching events plotted by SCFA treatment over time. Each dot indicates the time of an observed switching event switching event. **E** Mean ± SEM probability of cell-type switching during each growth phase by treatment. Two-way ANOVA followed by Tukey’s Honest Significance test, ** p < 0.001 **F** Results from Gillespie simulations using our experimentally measured parameters. Average of 100 individual simulations per condition. One-way ANOVA, ** p < 0.001.

Single-cell growth rates of untreated cells changed over time. Growth rates increased as they exited lag phase and entered exponential growth and decreased as they entered late exponential phase (Fig 3B, *two-way anova, p < 0.015 for all timepoints*). All cells continued growth into stationary phase (Fig S2). At all timepoints, SPI-1− cells grew approximately 25% faster than SPI− 1+ cells, in line with previous reports (Fig 3B,C, *two-way anova, p < 0.001,* (24)).

Cell-type switching rates of untreated cells also changed by growth phase (Fig 3D,E, *anova, p < 0.001*). Consistent with few SPI-1+ cells during pre-exponential phase growth (Fig 1E), SPI-1+ to SPI-1− switches were frequent during lag phase. Upon reaching early and late exponential phase, the rate of SPI-1− to SPI-1+ switches increased and SPI-1+ to SPI-1− switches decreased.

*Salmonella* cells grown in the presence of SCFAs behaved differently. Individual cells grown in the presence of SCFAs grew more slowly during lag phase, in line with our population-level observations (Fig 1A,B, S1). During the later phases of growth, SCFA treatment slowed single-cell growth rates by both cell-types during each growth phase, however to different degrees (Figs 3B,C). While SCFAs resulted in SPI-1− single cell growth rates approximately 25% lower than untreated cells, SCFAs slowed the growth of SPI-1+ cells by roughly 50% (Fig 3B). During each phase of growth, SCFA treated SPI-1+ cells grew roughly 50% slower than SPI-1− cells (Fig 3C).

Given that SCFAs were expected to repress SPI-1 expression, it was surprising to see that this was not the case (Fig 3D,E, Fig S2, Supplemental movie I). Rather, SPI-1+ cells frequently maintained expression for many generations, and switching off events were only observed during lag phase (Fig 3D,E). SPI-1− cells seldom switched to SPI-I+ in the presence of SCFAs, although these events were observed more frequently compared to SPI-1+ to SPI-1− switching during exponential growth. Although SCFAs reduced the number of cell-type switching events overall, the difference was not statistically significant (Fig 3D,E, *anova, p > 0.5*).

All of the measured changes in parameters can help to explain how SCFAs lower the frequency of SPI-1+ cells. To understand the interplay of these parameters, we used stochastic simulations to predict the proportion of SPI-1+ cells in a population growing under experimentally measured single-cell growth and switching values. Simulations using parameters extracted from SCFA untreated and treated cells yielded a final SPI-1+ frequency of 18% and 6%, respectively (Fig 3F). Both are roughly 2-fold lower than what is observed experimentally. This underestimation by the simulations suggests additional parameters are necessary to fully simulate SPI-1 subpopulation dynamics (expanded upon in Supplemental Materials). Nonetheless, these simple simulations captured the SCFA-mediated reduction SPI-1+ frequency in the population observed during our experiments.

These simulations can also inform on the relative influence that experimentally measured parameter shifts have on the proportion of SPI-1+ cells in a population. Populations of cells simulated to grow at untreated rates and cell-type switching at SCFA rates yielded a final SPI-1+ proportion similar to untreated cells (Fig 3F, *anova, p > 0.02).* Similarly, populations simulated to grow at SCFA-treated rates and switch with untreated rates more resembled SCFA-treated SPI-1+ proportions. These simulations indicate growth rate changes during SCFA treatment is a dominant driver of decreased SPI-1+ population frequency.

These results demonstrate SCFAs decrease SPI-1 expression at the population-level by acting on single-cell growth rates. We conclude SPI-1 expression is not inhibited by SCFAs as cells maintain SPI-1 expression in their presence for at least 12 h and over many generations. Taken together, our results indicate SCFAs decrease the proportion of SPI-1+ cells predominately through a selectively stronger reduction in growth by SPI-1+ cells.

### SPI-1+ cells maintain a higher PMF that is susceptible to perturbation by SCFAs

SCFAs can impact *Salmonella* physiology in multiple ways. *Salmonella* can use all three SCFAs present in our experiments as a carbon source, thus SCFAs could facilitate growth. More fundamentally, upon entry into the bacterial cytosol, SCFAs dissociate into the anion and a proton due to their relatively high pKa. An accumulation of the SCFA anion leads to an increase in turgor pressure while the accumulation of protons leads to a decrease in the intracellular pH (reviewed in (25)). A decrease in the intracellular pH ultimately lowers the proton motive force (PMF), leading to a slower rate of ATP production and by implication limited growth.

Because SCFAs slow single-cell growth rates, we examined how a global decrease in PMF impacts SPI-1+ population frequency. We added CCCP, a chemical decoupler of the proton gradient, at concentrations that did not inhibit population-level growth (Fig S4, *anova, p > 0.2*). Similar to SCFA treatment, the addition of CCCP decreased population-level SPI-1 expression (Fig 4A, *anova, p < 0.001*) and the proportion of SPI-1+ cells (Fig 4B, *anova followed by Tukey’s Honest Test, no SCFAs vs CCCP, p < 0.0001; + SCFAs vs CCCP, p > 0.5*). Thus, PMF dissipation at levels that do not limit population-level growth are sufficient to decrease the SPI-1+ subpopulation, consistent with the idea that SCFAs reduce the proportion of SPI-1+ cells through PMF dissipation.

**Fig 4:**
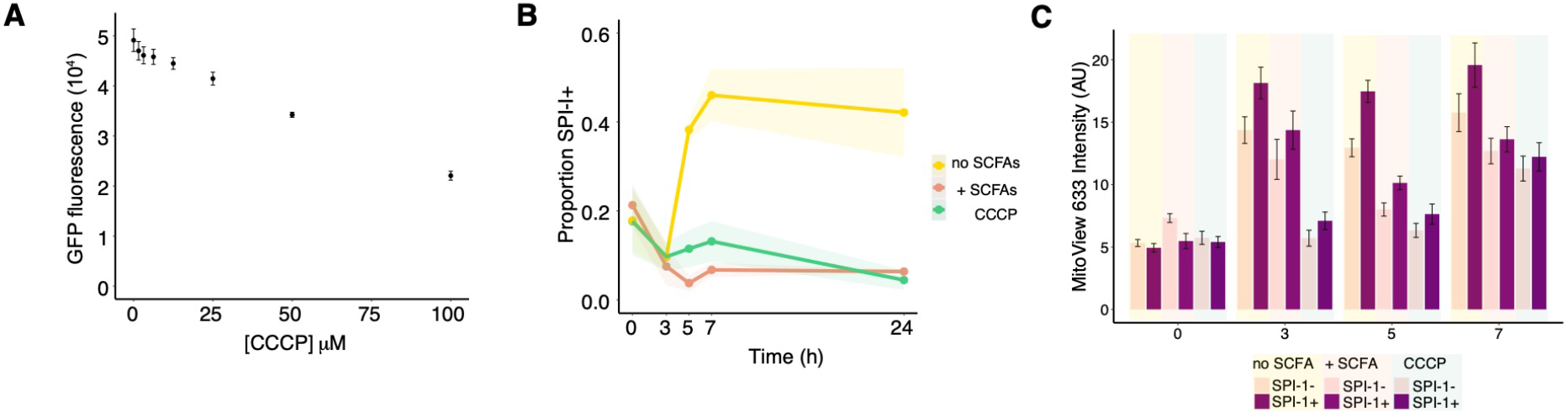
SCFAs and CCCP similarly impact SPI-1+ cell-type proportion and PMF dissipation. **A** Mean ± SEM maximal population-level SPI-1 expression by CCCP concentrations. Average of 3 independent experiments with triplicate wells; one-way ANOVA, p < 0.001. **B** Mean ± SEM SPI-1+ proportions over time in untreated, 150 mM SCFA, or 100 μM CCCP as determined by flow cytometry. Average of 4 independent experiments, 50,000 cells per sample per timepoint; two-way ANOVA followed by Tukey’s Honest Significance Test, ** p < 0.001. **C** Mean ± SEM membrane potential of SPI-1 cell-types by treatment as determined by MitoView 633 staining combined with flow cytometry. Average of 4 independent experiments. Three-way ANOVAs followed by Tukey’s Honest Significance Test, SPI-1− vs SPI-1+, p < 0.025; no SCFA vs + SCFA, p < 0.005; no SCFA vs CCCP, p < 0.001; no SCFA SPI-1+ vs + SCFA SPI-1+, p < 0.05; no SCFA SPI-1− vs + SCFA SPI.I−, p > 0.2; no SCFA SPI-1+ vs CCCP SPI-I+, p < 0.0005; no SCFA SPI-1− vs CCCP SPI-1−, p < 0.05; SCFA SPI-1+ vs CCCP SPI-1+, p > 0.5; SCFA SPI-1− vs CCCP SPI-1−, p > 0.4.

A remaining question is how SCFAs differentially affect the growth rates of SPI-1+ and SPI-1− cells. By combining single-cell analyses with a fluorescent PMF indicator, we observed SPI-1+ cells maintain a higher membrane potential compared to SPI-1− cells (Fig 4C, *three-way ANOVA, p < 0.05*). SCFA and CCCP treatment lead to an overall decrease in membrane potential (Figs 5A, *three-way anova, p < 0.005*). In particular, SCFA treatment led to a statistically significant decrease in the SPI-1+ cell PMF (*three-way anova followed by Tukey’s Honest Significance Test, p < 0.05),* whereas the SCFA-mediated decrease in in the SPI-1− cell PMF was not statistically significant (*p > 0.2; no interaction detected*).

Collectively, these observations are in line with the idea that SCFAs preferentially slow SPI-1+ cell growth by decreasing their PMF to a higher degree. We suspect SPI-1+ cells are more sensitive to SCFA-mediated PMF reduction due to their inherently higher PMF.

### Conclusions

Our study demonstrates SCFAs decrease population-level SPI-1 expression by differentially impacting SPI-1 cell-type behaviors. Although not the major driver, we find SCFAs influence the molecular regulation of SPI-1 by reducing cell-type switching events. However, because SCFA-treated SPI-1+ cells maintain expression across many generations and SPI-1− cells initiate SPI-1 expression, we conclude SCFAs do not act as a canonical transcriptional co-repressor of SPI-1 transcription. Our data collectively indicate SCFAs shape population-level SPI-1 expression predominately by acting on the growth rate of SPI-1+ cells.

These findings also have implications for how *Salmonella* causes disease. That SCFAs act on cell-type proportions indicates they shape the cooperative interactions between SPI-1− and SPI-1+ subpopulations during infection (15–17, 26). Because SCFA levels fluctuate as a function of nutrient intake, microbiota metabolism, and location in the gut, these dynamic conditions likely influence on *Salmonella* cell-type interactions over time. High SCFA levels would lead to fewer inflammation-inducing SPI-1+ cells and thus limit the expansion of SPI-1− cells, while low SCFA levels would enhance inflammation induction by increasing the number of SPI-1+ cells and therefore aid in *Salmonella* colonization of the gut. SCFA levels and their dynamics, therefore, likely help to balance infection outcomes between colonization resistance, asymptomatic carriage, and symptomatic disease.

Lastly, there is strong support for the idea that SPI-1+ and SPI-1− cell-types are distinct in ways other than SPI-1 expression. Previous reports show these two cell-types have different growth rates, susceptibility to antibiotics, and cell sizes (17, 27). We add to this body of evidence of *Salmonella* differentiation by showing that SPI-1+ and SPI-1− cells differ in response to an environmental stimulus (SCFAs) and PMF. We speculate these reflect further specialization by each cell type to best fulfill their function. For example, the higher PMF of SPI-1+ cells could allow for more efficient effector secretion during interaction with host cells, whereas the maintenance of a high PMF by SPI-1− cells would be unconducive for fast growth (28). Identifying distinguishing characteristics of the two cell-types will shed light on the specializations each make to fulfill their function, including how SPI-1+ cells prepare for their intracellular future.

As single-cell analyses have become more common, the importance of bacterial cell-types during pathogenesis is becoming clear (15–17, 29–34). Combining these efforts with developing detailed molecular maps of each cell-type will help guide the development of therapeutics which inhibit a single cell-type of interest (*e.g.* virulent cells) rather than the whole population. Cell-type targeted therapeutics could impede disease development while not selecting against a given species as only a subset of the cells would be susceptible (35, 36), a valuable strategy in the fight against antibiotic resistance.

## Materials and Methods

### Bacterial cultivation

Throughout the study we used SB300 (a spontaneous Streptomycin resistant SL1344 derivative, wild-type (wt)) and its SPI-1 fluorescent transcriptional reporter, P*prgH-gfp* (13). Frozen strains were streaked onto LB Miller agar for single colonies. After 24 h, a single colony was transferred to 5 mL LB Miller in a 15 mL round bottom tube and incubated at 37 °C with shaking at 200 rpm. A 16-18 h liquid culture prepared in this way was the starting point for all experiments. All experiments were performed in LB Miller and, when applicable, supplemented with sodium acetate, sodium butyrate, and sodium propionate at appropriate concentrations (table S1).

### Quantification of population-level growth and SPI-1 expression

We prepared a 96-well plate containing 2 μL of an overnight in 200 μL of media containing or not SCFAs at indicated concentrations. OD600 and GFP fluorescence of cultures were measured every 3 minutes using a heated, automated microplate reader (Biotek Synergy). Three experiments were performed with each condition in triplicate. Blank (uninoculated) and autofluorescence (SB300) background control wells were run in each experiment. Resultant values were background subtracted by appropriate time-matched, control-well values. Maximum growth rate, lag time, yield, and max GFP fluorescence levels were determined by fitting a logistic equation to each time series using the GrowthCurver package in R (37).

### Single-cell SPI-1 expression quantification using flow cytometry

Overnight cultures of wt and SPI-1 reporter cells were diluted 1:100 into LB Miller ± SCFAs and incubated at 37°C with shaking. At each timepoint, 10 mL of the culture was centrifuged (tabletop centrifuge,3000 *xg*, 15 mins). Supernates were removed and pellets were resuspended in 1 mL PBS. Cells were centrifuged (microcentrifuge, 14000 *xg*, 5 mins), pellets resuspended in 1 mL 4% paraformaldehyde in PBS, and incubated at room temperature for 10 min. Fixed cells were pelleted, washed in 1 mL PBS, resuspended in 1 mL PBS, and stored at 4°C. Fixed cell samples were analyzed by flow cytometry within 1 week of collection. Forward scatter, side scatter, and GFP fluorescence of 50,000 events from each sample were measured using a Beckman Coulter Gallios flow cytometer. We performed three independent experiments.

Average per-cell gene expression (mean fluorescence intensity, MFI) and subpopulation frequencies were calculated as follows. Distributions of GFP fluorescence values per sample were extracted from .fcs files using FlowCore (38). A test for bimodality was performed on each log10 transformed GFP fluorescence distribution (39). If distributions tested negativity for bimodality (*e.g.* all wt samples), a single normal distribution was fitted to the data. If distributions tested positively for bimodality (almost all reporter strain samples), the mixtools package was used to fit two normal distributions to the bimodal distribution (40). The mean of each distribution is the MFI of the subpopulation and the proportion of events assigned to each distribution using mixtools was used as the subpopulation frequency. The MFI of wt samples was used as a benchmark to determine which MFI represented the SPI-1− subpopulation. There were no significant differences between wt and SPI-1− reporter cell MFIs.

### Mathematical modeling and simulations

The simple differential equation was solved using Mathematica. Analytical solutions were run in MatLab. Gillespie simulations were run in RStudio. For more information, please see Supporting Materials.

### Feeding-culture microfluidics

Microfluidic chips were fabricated using previously described methods (23, 27). All images were acquired with an automated Olympus IX81 inverted microscope using an oil-immersion 100x objective (Olympus), an ORCA-flash 4.0 v2 sCMOS camera (Hamamatsu), X-Cite120 metal halide arc lab (Lumen Dynamics), and Chroma 49000 fluorescent filter sets (Chroma, N49002).

A 16 – 18 h stationary culture of P*prgH-gfp* reporter cells were loaded into the microfluidic device. Cells were concentrated for loading into the chip by centrifuging 100 μL of the stationary phase culture, decanting excess media, and resuspending the cell pellet in the remaining culture media. One microliter of 1% Tween-20 in PBS was added to facilitate loading cells into observation channels. The concentrated cells were then pipetted into the microfluidic device and examined by microscopy for sufficient loading. We then connected flasks of LB or LB + SCFAs (150 mM) to the microfluidic chip to feed the cells with a flow rate of 0.5 mL per hour.

To recapitulate the population-level experiments above, we used a feeding culture approach. After fresh media was flowed through the device for approximately 0.5-1 h (time used to set-up the automated, time-lapse program for image acquisition), we inoculated the flasks feeding the microfluidic chip with stationary phase reporter cells at a ratio of 1:100. We then started image acquisition (t=0). Phase contrast and GFP images were acquired for each position at 3-minute intervals over 12 hours.

### Calculation of growth rates and cell-type switching probabilities

Time-lapse microscopy images were segmented, tracked, and quantified using SuperSegger (41). Images were first deconvolved using a point-spread function (42). Segmentation of images (identification of individual cells) was performed using optimized segmentation constants to detect both SPI-1+ and SPI-1− cell types, which differ in size and curvature. All segmentation and tracking results were manually curated for erroneous boundary-calling and tracking. For all lineages examined, we only tracked and quantified the “mother cell,” the cell at the closed end of the observation channel.

Cell-type switches were called by examining GFP traces of each mother cell over time. Mean GFP values per cell over the duration of the experiment were smoothed by fitting a LOESS curve (α = 0.5) to reduce noise. The first derivative of the smoothed GFP trace was then examined to determine changes in SPI-1 expression: values of approximately zero (±10, MFI change of 10 GFP AU over 30 minutes) indicates equal GFP levels over time; values > 10 indicate increases in MFI levels over time; and values < −10 indicate decreases in MFI levels over time. The time in which the derivative surpassed 10 or dropped below −10 was considered the time of cell-type switch.

The time and direction of cell-type switch was then mapped to individual cells to call which cell-type each cell was in at a given time. Manual inspection indicated good agreement between individual cell MFI levels and cell-type mapping using the above calling procedure. Growth rates of individual cells by cell-type assignment were then plotted and examined statistically using two-way analysis of variance.

### CCCP treatment

Reporter cells in a range of concentrations of CCCP were analyzed by 96-well plate assays as described above. The flow cytometric analyses were performed and analyzed as described above by adding 100 μM CCCP to the culture medium.

### Single-cell PMF measurements

Reporter cells were grown in LB, LB + 150 mM SCFAs, LB + 150 mM NaCl (ionic control for SCFAs), LB + 100 μM CCCP, and LB + DMSO (CCCP vehicle control) and sampled at 0, 3, 5, and 7 hours post-inoculation. At the time of sampling, cells were centrifuged to pellet (table-top centrifuge, 3000 *xg*, 15 min) and resuspended in PBS containing MitoView 633 (Biotium). MitoView 633 accumulates in the inner membrane as a function of membrane potential. Higher fluorescence signal indicates higher membrane potential, i.e. a higher charge differential across the membrane. Sampled reporter cells were stained with MitoView-633 for 20 minutes on ice and then analyzed using flow cytometry to assign SPI-1 cell-type and MFI of MitoView accumulation. Wt *Salmonella* cells (no reporter) were used as a negative control to confirm no bleed-through of GFP or MitoView into the other fluorescence channels. All MitoView values were background subtracted with the time-matched, unstained, reporter strain control. At all timepoints, the addition of 150 mM NaCl and DMSO lead to an approximate 20% increase in MitoView accumulation; an expected observation as the addition of ions will impact the charge differential across the membrane. Thus, a 20% correction was applied to SCFA-treated and CCCP-treated cells to account for the impact of 150 mM extra ions or DMSO in the environment.

## Acknowledgments

The authors would like to express gratitude to all members of the Microbial Systems Ecology group for the helpful discussions during all phases of this work.

---

**Fig S1:**
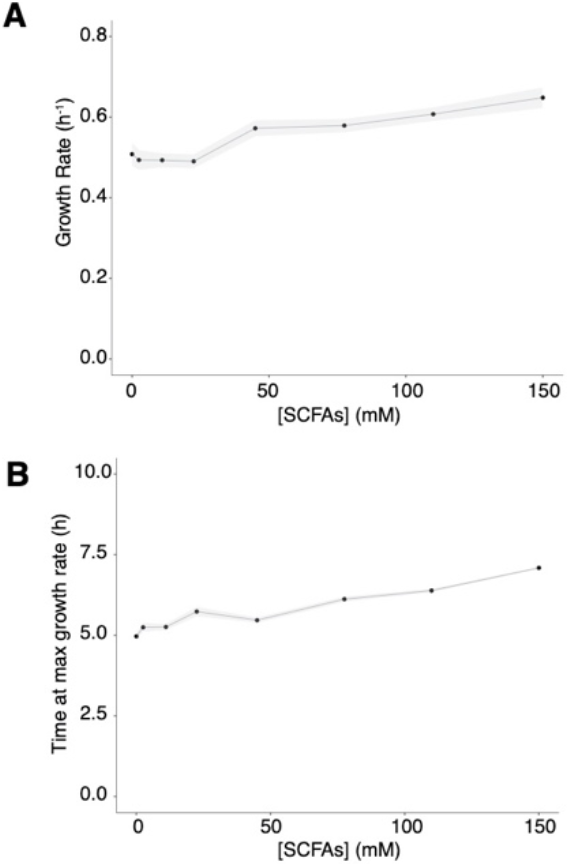
Impact of SCFAs on population-level growth. **A** SCFA treatment correlates with higher maximal population-level growth, one-way ANOVA, p <0.005 **B** SCFA treatment delays time to maximal growth rate, a proxy for lag-time, one-way ANOVA, p < 0.0001

**Fig S2:**
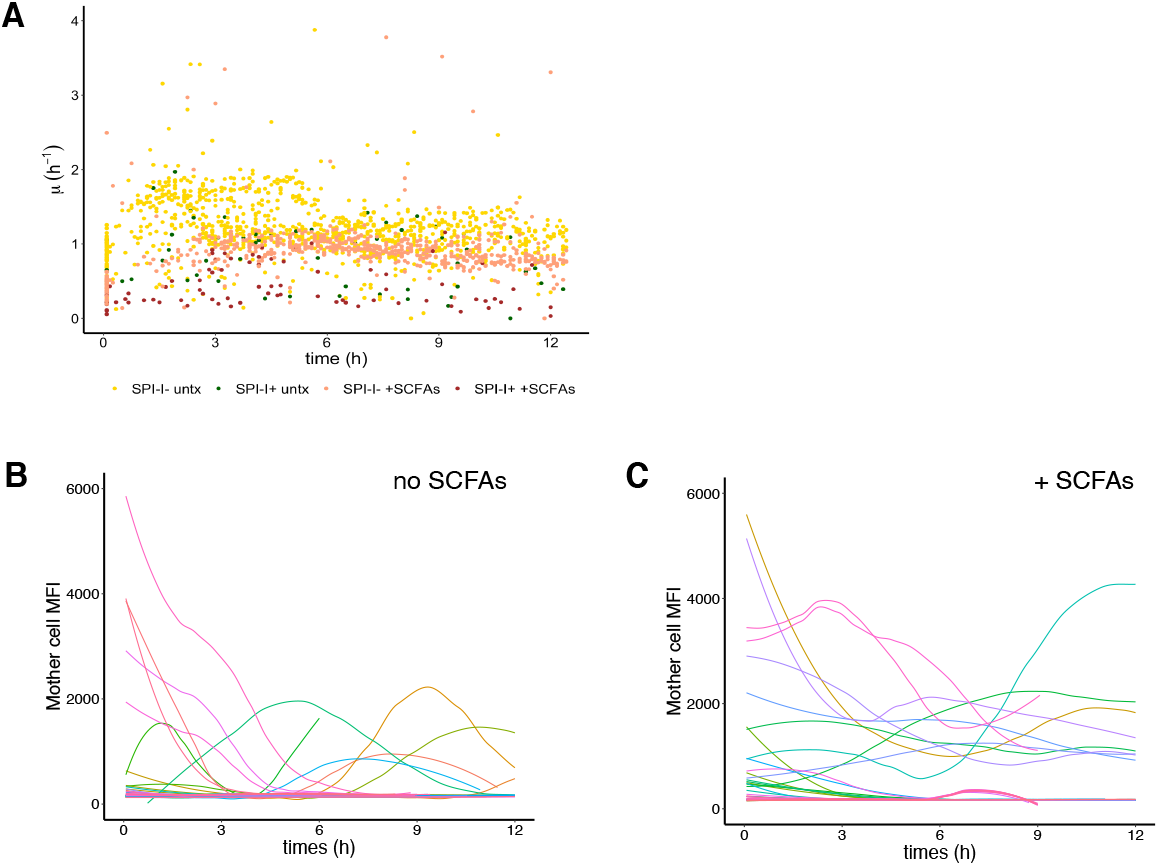
Single-cell growth rates and SPI-1 expression over time. **A** Single-cell growth rate of every cell included in our analysis by time. SCFA treatment and SPI-1 cell-type indicated by color. Mean GFP intensity (SPI-1 expression) of lineages of cells over time in **B** untreated and **C** SCFA-treated conditions. In panels B and C, flat lines at close to zero indicate SPI-1− cells.

**Fig S3:**
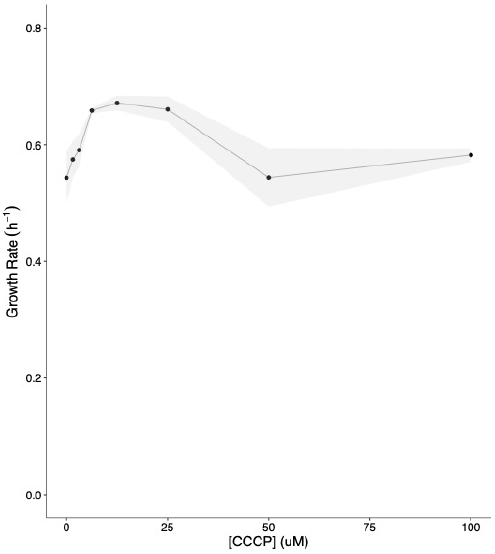
Impact of CCCP on population-level growth. Mean ± SEM population-level growth rate by CCCP treatment. Average of 3 independent experiments with triplicate wells; one-way ANOVA, p < 0.001)

**Supplemental video 1: Population-level vs single-cell growth and SPI-1 expression by SCFA treatment**

**Population-level** growth (blue) and SPI-1 expression by reporter cells with or without 150 mM SCFAs as quantified in our plate reader experiments (Fig 1). **Single-cell** growth and SPI-1 expression from our time-lapse microscopy experiments in microfluidic devices (Fig 3). Movies of population-level and single-cells are synced in time.

**Table S1:** SCFA concentrations used.

| **Total SCFAs (mM)** | **Sodium Acetate (mM)** | **Sodium Butyrate (mM)** | **Sodium Propionate (mM)** |
| --- | --- | --- | --- |
| **0** | 0 | 0 | 0 |
| **2.5** | 1 | 1 | 0.5 |
| **11** | 5 | 5 | 1 |
| **22.5** | 10 | 10 | 2.5 |
| **45** | 25 | 15 | 5 |
| **75** | 50 | 18 | 7 |
| **110** | 75 | 25 | 10 |
| **150** | 100 | 25 | 25 |

## Supporting Materials

## 1 Mathematical model

Here, we present the details of the mathematical model to predict SPI-1+ and SPI-1− cell-type frequencies as a function of growth and cell-type switching rates shown in Figure 2 of the main text. This model di↵ers from the model previously described by Sturm, *et al*, (2011) by including SPI-1+ to SPI-1− switching events, as these are observed experimentally. In addition, we discuss cases for specific choice of the parameter space.

### 1.1 Two growth rates and two switching rates model

SPI-1− and SPI-1+ cell number is denoted with *X* and *Y*, respectively. SPI-1− cells grow with growth rate *μ*_*X*_ and switch to SPI-1+ cells with rate *δ*_*X→Y*_ = *δ*_*YX*_. SPI-1+ cells grow with growth rate *μ*_*Y*_ and switch to SPI-1− cells with rate *δ*_*Y→X*_ = *δ*_*XY*_. This models exponential growth of the two cell-types assuming no interactions other than cell-type switching.

The quantity of each cell-type follows this system of ordinary differential equations:

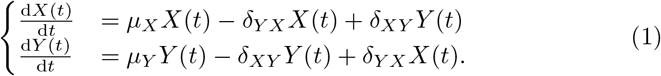

The solution of the system of differential equations is:

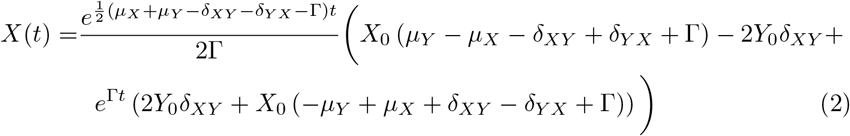

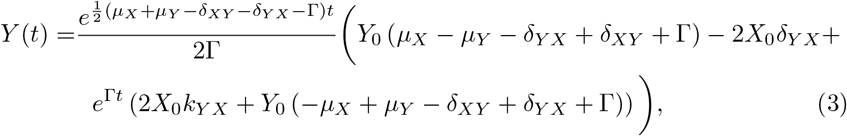

where *X*_0_ and *Y*_0_ are the initial number of cells at time zero, and

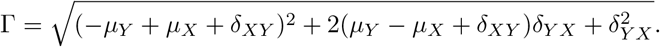

The fraction of SPI-1+ cells in a population growing exponentially, for time *t* ≫ Γ > 0 reach a steady state at

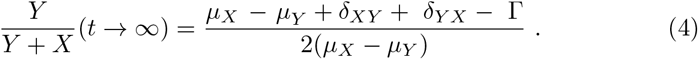

The fraction of SPI-1+ cells in Eq. (4) is shown in Figure 2 of the main text as a function of the growth rates (Fig. 2c) and switching rates (Fig. 2d).

### 1.2 The case of negligible switching from SPI-1+ to SPI-1−: *δ*_*YX*_ =0

For simplicity, we describe here the case in which the switching rate from SPI-1+ to SPI-1− is negligible *k*_*YX*_ = 0, and the growth rate of the SPI-1− cells is larger than SPI-1+ cells, *μ*_*X*_ > *μ*_*Y*_. In this case, the solution of the simplify version of Eq. (1) are:

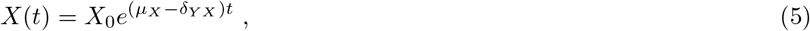

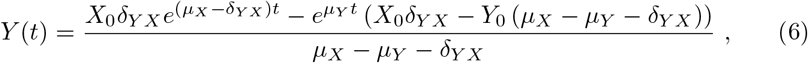

The fraction of SPI-1+ cells in a population growing exponentially, under the assumptions that *μ*_*X*_ > *μ*_*Y*_ + *δ*_*YX*_ reach a steady state at

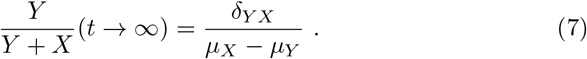

### 1.3 The case of equivalent growth rates for SPI-1+ and SPI-1−: *μ* = *μ*_*X*_ = *μ*_*Y*_

The case of equivalent growth rates between the two cell-types has been use to resolve the indeterminacy of the solution of Eq.(4) for *μ*_*X*_ = *μ*_*Y*_ = *μ*. In this case, the solution of the version of Eq. (1) are:

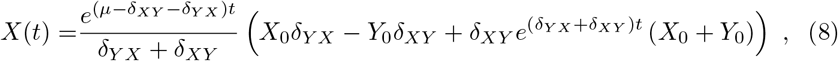

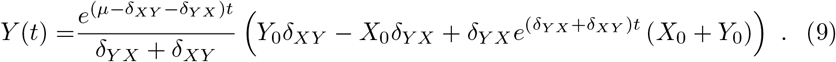

The fraction of SPI-1+ cells in a population growing exponentially reach a steady state at

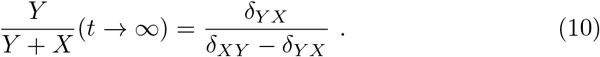

## 2 Growth Simulations

In this section, we outline the details of the growth simulations presented in Figure 3E of the main text.

### 2.1 Rationale

The analytical solution presented above models populations growing exponentially, however cells in our experiments grow logistically, plateauing upon approaching the carrying capacity. To better describe our experimental system where cells grow logistically and cell-type switches are stochastic, we moved to stochastic simulations of cell-type frequencies using the parameters of growth, cell-type switching, and carrying capacity.

### 2.2 Growth, switching, and carrying capacity simulations

We used Gillespie stochastic simulations to estimate cell-type proportions during logistic growth. At each randomly determined time interval, one of four possible events can occur:

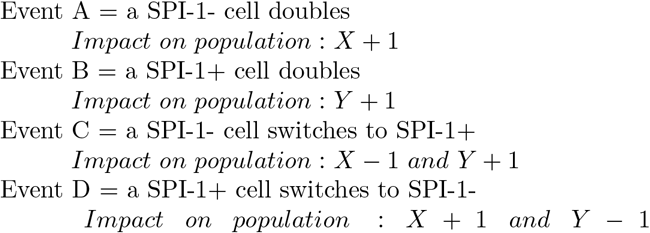

A random number from a uniform distribution between 0 and 1 was selected at each iteration to determine the outcome of the event (*X* + 1, etc). For example, if the random number was greater than *P*_*A*_ and less than *P*_*B*_, the outcome would be *Y* + 1. The probabilities were as follows, where *k* is the carrying capacity:

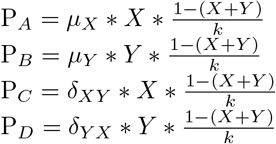

Time elapsed between events during our simulations was determined as follows:

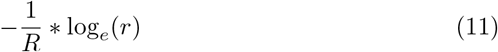

Where *R* is the sum of probabilities and *r* is a randomly selected number from a uniform distribution between 0 and 1.

### 2.3 Parameters used

For these simulations, we used the mean of experimentally measured parameters for single-cell growth and cell-type switching rates during lag (1), early exponential (2), and late exponential (3) growth phases. This was done to account for the different cell-type switching rates during the different growth phases and to also recapitulate the delayed growth during lag phase not accounted for in the base equation. Growth was simulated for 12 hours.

For all simulations, *X*_*o*_ = 60, *Y*_*o*_ = 40, and *k* = 10000, approximating the 1:100 culture dilution we performed during the experiment. Lag phase values were used for the first 30 minutes of the simulation; early exponential for the next 6 h; and late exponential for the last 5.5 h.

#### 2.3.1 No SCFAs

All values are provided in events per hour.

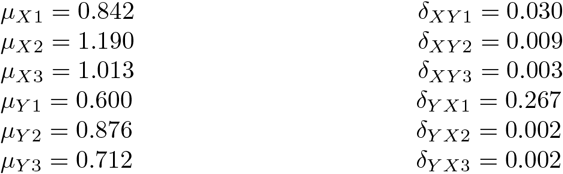

#### 2.3.2 + SCFAs

All values are provided in events per hour.

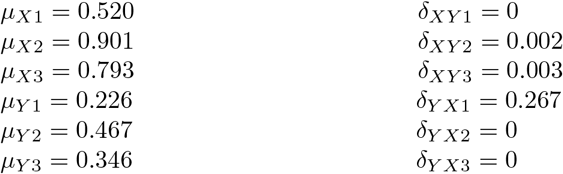

### 2.4 Potential reasons for underestimation of SPI-1+ cell proportion during simulations

While our simulations using parameters quantified by time-lapse microscopy captured the SCFA-mediated decrease in SPI-1+ proportion, they underestimated the experimentally measured SPI-1+ proportion by roughly 2-fold in both the absence and presence of SCFAs. Here, we speculate on why this may be the case.

#### 2.4.1 Assumptions in the simulation

For each switching event, we simulate a single cell turning on at a time (*X* − 1, *Y* + 1). By observing real switching events, we see that 2-4 sister cells (mean = 3.111 cells/switch event, see Supplementary video) simultaneously turn from SPI-1- to SPI-1+ (*X −* 3.111, *Y* + 3.111). When we attempt to account for this in our simulations, we observe a 3% and 0.8% increase in the SPI-1+ cell-type frequency using untreated and SCFA parameters, respectively. This coordinated switching by sister cells does therefore impact the final proportion, but does not account for all of the underestimation.

The only parameters we use here are growth rates of SPI-1- and SPI-1+ cells and cell-type switching rates. It’s likely additional parameters are necessary to better predict cell-type frequencies. For example, we model interactions between the cell types only in the carrying capacity and switching rates; other types of interactions, including chemical communication, are neglected.

#### 2.4.2 Potential biological sources of noise

As mentioned above, during experiments, on average 3 cells turn on at a time. This suggests the signal for cell-type switching takes place in a cell’s “mother” or “grandmother” and then executed 1-2 generations later. This suggests there are influences other than only single-cell growth rates and switching frequencies on SPI-1+ proportion, including history dependence in SPI-1− to SPI-1+ switches. This makes it very difficult to simulate with precision. This aspect is currently under investigation. As an aside, during switch off events, we seldom observe sister cells turning off SPI-1 expression simultaneously, indicating a lack of history dependence for SPI-1+ to SPI-1− events.

Cells which have just switched to the SPI-1+ phenotype also undergo a reductive division soon after becoming GFP+, doubling the SPI-1+ cell-type soon after switches. This dynamic is also very difficult to capture using stochastic simulations and could be a source of the underestimation.

Although we employed a feeding-culture approach, cells in the microfluidic chip, from where our single-cell parameters were derived, are experiencing a different environment compared to cells growing in liquid culture as in our population-level experiments (e.g. physical constraint and likely differences in oxygen levels as PDMS is very permeable to gasses). We also therefore cannot exclude the difference in physico-chemical environment as be a source of error in parameter estimation.

## References

1. J. H. Cummings, E. W. Pomare, H. W. J. Branch, E. Naylor, G. T. Macfarlane, Short chain fatty acids in human large intestine, portal, hepatic and venous blood. 28, 122–1 (1987).

2. L. M. Filkins, et al., Prevalence of Streptococci and Increased Polymicrobial Diversity Associated with Cystic Fibrosis Patient Stability. J. Bacteriol. 194, 4709–4717 (2012).

3. D. Ríos-Covián, et al., Intestinal Short Chain Fatty Acids and their Link with Diet and Human Health. Front. Microbiol. 7, 185 (2016).

4. E. G. Zoetendal, et al., The human small intestinal microbiota is driven by rapid uptake and conversion of simple carbohydrates. ISME J. 6, 1415–1426 (2012).

5. D. L. LaRock, A. Chaudhary, S. I. Miller, Salmonellae interactions with host processes. Nat. Rev. Microbiol. 13, 191–205 (2015).

6. J. E. Galán, R. Curtiss, Cloning and molecular characterization of genes whose products allow Salmonella typhimurium to penetrate tissue culture cells. Proc. Natl. Acad. Sci. U. S. A. 86, 6383–7 (1989).

7. S. Y. Wotzka, B. D. Nguyen, W.-D. Hardt, Salmonella Typhimurium Diarrhea Reveals Basic Principles of Enteropathogen Infection and Disease-Promoted DNA Exchange. Cell Host Microbe 21, 443–454 (2017).

8. P. A. McLaughlin, et al., Inflammatory monocytes provide a niche for Salmonella expansion in the lumen of the inflamed intestine. PLOS Pathog. 15, e1007847 (2019).

9. B. Stecher, et al., Salmonella enterica Serovar Typhimurium Exploits Inflammation to Compete with the Intestinal Microbiota. PLoS Biol. 5, e244 (2007).

10. S. E. Winter, et al., Gut inflammation provides a respiratory electron acceptor for Salmonella. Nature 467, 426–9 (2010).

11. P. Thiennimitr, et al., Intestinal inflammation allows Salmonella to use ethanolamine to compete with the microbiota. Proc. Natl. Acad. Sci. 108, 17480–17485 (2011).

12. L. Maier, et al., Microbiota-derived hydrogen fuels Salmonella typhimurium invasion of the gut ecosystem. Cell Host Microbe 14, 641–51 (2013).

13. I. Hautefort, M. J. Proença, J. C. D. Hinton, Single-copy green fluorescent protein gene fusions allow accurate measurement of Salmonella gene expression in vitro and during infection of mammalian cells. Appl. Environ. Microbiol. 69, 7480–91 (2003).

14. D. Bumann, Examination of Salmonella gene expression in an infected mammalian host using the green fluorescent protein and two-colour flow cytometry. Mol. Microbiol. 43, 1269–1283 (2002).

15. M. Diard, et al., Stabilization of cooperative virulence by the expression of an avirulent phenotype. Nature 494, 353–356 (2013).

16. M. Ackermann, et al., Self-destructive cooperation mediated by phenotypic noise. Nature 454, 987–990 (2008).

17. A. Sturm, et al., The Cost of Virulence: Retarded Growth of Salmonella Typhimurium Cells Expressing Type III Secretion System 1. PLoS Pathog. 7, e1002143 (2011).

18. S. Y. Wotzka, et al., Escherichia coli limits Salmonella Typhimurium infections after diet shifts and fat-mediated microbiota perturbation in mice https://doi.org/10.1038/s41564-019-0568-5 (February 10, 2021).

19. S. D. Lawhon, R. Maurer, M. Suyemoto, C. Altier, Intestinal short-chain fatty acids alter Salmonella typhimurium invasion gene expression and virulence through BarA/SirA. Mol. Microbiol. 46, 1451–1464 (2002).

20. I. Gantois, et al., Butyrate specifically down-regulates salmonella pathogenicity island 1 gene expression. Appl. Environ. Microbiol. 72, 946–9 (2006).

21. F. Rivera-Chávez, et al., Depletion of Butyrate-Producing Clostridia from the Gut Microbiota Drives an Aerobic Luminal Expansion of Salmonella. Cell Host Microbe 19, 443–454 (2016).

22. A. Jacobson, et al., A Gut Commensal-Produced Metabolite Mediates Colonization Resistance to Salmonella Infection. Cell Host Microbe 24, 296–307.e7 (2018).

23. S. Moreno-Gámez, et al., Wide lag time distributions break a trade-off between reproduction and survival in bacteria. Proc. Natl. Acad. Sci. U. S. A. 117, 18729–18736 (2020).

24. A. Sturm, et al., The Cost of Virulence: Retarded Growth of Salmonella Typhimurium Cells Expressing Type III Secretion System 1. PLoS Pathog. 7, e1002143 (2011).

25. A. J. Wolfe, The acetate switch. Microbiol. Mol. Biol. Rev. 69, 12–50 (2005).

26. M. A. Sánchez-Romero, J. Casadesús, Contribution of SPI-1 bistability to Salmonella enterica cooperative virulence: insights from single cell analysis. Sci. Rep. 8, 14875 (2018).

27. M. Arnoldini, et al., Bistable Expression of Virulence Genes in Salmonella Leads to the Formation of an Antibiotic-Tolerant Subpopulation. PLoS Biol. 12, e1001928 (2014).

28. M. Erhardt, M. E. Mertens, F. D. Fabiani, K. T. Hughes, ATPase-Independent Type-III Protein Secretion in Salmonella enterica. PLoS Genet. 10, e1004800 (2014).

29. K. M. Davis, S. Mohammadi, R. R. Isberg, Community behavior and spatial regulation within a bacterial microcolony in deep tissue sites serves to protect against host attack. Cell Host Microbe 17, 21–31 (2015).

30. K. M. Davis, R. R. Isberg, Defining heterogeneity within bacterial populations via single cell approaches. BioEssays 38, 782–790 (2016).

31. K. M. Davis, For the Greater (Bacterial) Good: Heterogeneous Expression of Energetically Costly Virulence Factors. Infect. Immun. 88(2020).

32. E. H. Rego, R. E. Audette, E. J. Rubin, Deletion of a mycobacterial divisome factor collapses single-cell phenotypic heterogeneity. Nature 546, 153–157 (2017).

33. M. Diard, et al., Antibiotic Treatment Selects for Cooperative Virulence of Salmonella Typhimurium. Curr. Biol. 24, 2000–2005 (2014).

34. I. Ronin, et al., A long-term epigenetic memory switch controls bacterial virulence bimodality. Elife 6, 7808–7818 (2017).

35. D. A. Rasko, V. Sperandio, Anti-virulence strategies to combat bacteria-mediated disease. Nat. Rev. Drug Discov. 9, 117–128 (2010).

36. G. Bell, C. MacLean, The Search for “Evolution-Proof” Antibiotics. Trends Microbiol. 26, 471–483 (2018).

37. K. Sprouffske, A. Wagner, Growthcurver: an R package for obtaining interpretable metrics from microbial growth curves. BMC Bioinformatics 17, 172 (2016).

38. F. Hahne, et al., flowCore: a Bioconductor package for high throughput flow cytometry. BMC Bioinformatics 10, 106 (2009).

39. Martin Maechler, “Package ‘diptest’ Title Hartigan’s Dip Test Statistic for Unimodality-Corrected” (2016) (February 5, 2021).

40. T. Benaglia, D. Chauveau, D. R. Hunter, D. Young, **mixtools**: An *R* Package for Analyzing Finite Mixture Models. J. Stat. Softw. 32, 1–29 (2009).

41. S. Stylianidou, C. Brennan, S. B. Nissen, N. J. Kuwada, P. A. Wiggins, SuperSegger: robust image segmentation, analysis and lineage tracking of bacterial cells. Mol. Microbiol. 102, 690–700 (2016).

42. S. van Vliet, et al., Spatially Correlated Gene Expression in Bacterial Groups: The Role of Lineage History, Spatial Gradients, and Cell-Cell Interactions. Cell Syst. 6, 496–507.e6 (2018).

